# The rhesus monkey hippocampus critically contributes to scene memory retrieval, but not new learning

**DOI:** 10.1101/288407

**Authors:** Sean Froudist-Walsh, Philip G. F. Browning, Paula L. Croxson, Kathy L. Murphy, Jul Lea Shamy, Tess L. Veuthey, Charles R. E. Wilson, Mark G. Baxter

**Affiliations:** Glickenhaus Laboratory of Neuropsychology, Department of Neuroscience and Friedman Brain Institute, Icahn School of Medicine at Mount Sinai, New York, NY 10029, USA; Univ Lyon, Université Claude Bernard Lyon 1, Inserm, Stem Cell and Brain Research Institute U1208, 69500 Bron, France

**Keywords:** episodic, retrograde amnesia, anterograde amnesia, macaque, rhesus

## Abstract

Humans can recall a large number of memories years after the initial events. Patients with amnesia often have lesions to the hippocampus, but human lesions are imprecise, making it difficult to identify the anatomy underlying memory impairments. Rodent studies enable great precision in hippocampal manipulations, but not investigation of many interleaved memories. Thus it is not known how lesions restricted to the hippocampus affect the retrieval of multiple sequentially encoded memories. Furthermore, disagreement exists as to whether hippocampal inactivations lead to a temporally graded, or ungraded amnesia, which could be a consequence of differences between rodent and human studies. In the current study, four rhesus monkeys received bilateral neurotoxic lesions of the hippocampus, and were compared to thirteen unoperated controls on recognition and new learning of visual object-in-place scenes. Monkeys with hippocampal lesions were significantly impaired compared to controls at remembering scenes that were encoded before the lesion. We did not observe any temporal gradient effect of the lesion on memory recognition, with recent and remote memories being equally affected by the lesion. Monkeys with hippocampal lesions showed no deficits in learning and later recognising new scenes. Thus, the hippocampus, like other cortical regions, may be engaged in the acquisition and storage of new memories, but its role can be taken over by spared regions following a lesion. These findings illustrate the utility of experimental paradigms for studying retrograde and anterograde amnesia that make use of the capacity of nonhuman primates to rapidly acquire many distinct visual memories.

## Introduction

When memory works, we recall important details of experiences years later. Widely distributed patterns of brain activity are somehow consolidated, so that a record of activity that would otherwise be forgotten is kept. The brain mechanisms of such "systems consolidation" point towards a pivotal role for the hippocampus. For more than 60 years the hippocampus has been a focus of investigation for memory research, due to the devastating amnesia experienced by Henry Molaison following surgical resection of the hippocampus (Scoville and Milner, 1957). Reassessment of the original lesions demonstrated that the affected area was much greater than initially thought (Augustinack et al., 2014), complicating the interpretation of the role of the hippocampus, and other brain regions, in human episodic memory.

Considerable evidence on systems consolidation has come from studies in rodents using behavioral paradigms in which the time of learning is precisely defined, including inhibitory avoidance, contextual fear conditioning, and social transmission of food preference. These procedures are ideal for examining structural and biochemical changes minutes to weeks following learning, as well as reliance on different brain structures for retrieval at different times after learning. Many of these experiments have pointed towards the importance of the hippocampus in retrieval of recent memories, within 24 hours, and, conversely, the anterior cingulate cortex for more remote memories, about a month old (Frankland et al., 2004, 2006; Teixeira et al., 2006; Ding et al., 2008). Highly "schematized" memories that are variations of a familiar memory task may become hippocampal-independent much more rapidly, within hours (Tse et al., 2011). Multiple studies in humans have shown temporally limited retrograde amnesia after damage apparently limited to the hippocampus (Reed and Squire, 1998; Kapur and Brooks, 1999; Bayley et al., 2005). Nevertheless, some studies point toward a more enduring role for the hippocampus in memory retrieval. For example, humans that suffer transient global amnesia show focal lesions in hippocampal region CA1 and retrograde amnesia that spans decades (Bartsch and Deuschl, 2010; Bartsch et al., 2011). Conflicting evidence from human and rodent studies as to the temporal gradient of amnesia could reflect a dissociation between transient and sustained hippocampal inactivation (Goshen et al., 2011), while conflicting human studies may be a result of the lack of experimental control over the brain area affected. It is thus not clear whether complete permanent lesions restricted to the hippocampus will lead to temporally graded, or ungraded retrograde amnesia.

The hippocampus may coordinate neocortical activity rather than serve simply as a temporary memory store, as hippocampal inactivation impairs memory retrieval and reactivation of cortical neurons that were active at encoding (Tanaka et al., 2014) and direct reactivation of cortical neurons that were active at context encoding produces context-specific behavior even if the hippocampus is inactivated (Cowansage et al., 2014). Hippocampal memory traces may also remain active even while complementary memory traces become established in the cortex (Moscovitch et al., 2005; Tayler et al., 2013), suggesting that, in some cases hippocampal activation during memory retrieval may be commonplace, but not necessary. It is not clear whether the role of the hippocampus in coordination of cortical reactivation is only required if the hippocampus was also engaged during encoding of the memory, or whether hippocampal-independent mechanisms for coordination of cortical memory traces may also exist or develop following hippocampal lesions.

We sought to overcome some of the limitations of human and rodent memory studies by examining retrieval and new learning of visual object-in-place scenes in rhesus monkeys after selective, bilateral neurotoxic lesions limited to the hippocampus. Object-in-place scene memory is thought to closely model human episodic memory (Gaffan, 1994; Mitchell et al., 2008). Monkeys with hippocampal lesions were significantly impaired at remembering scenes that were encoded before the lesion, but showed no deficits in learning and later recognizing new scenes. We did not observe any temporal gradient effect of the lesion on memory recognition, with recent and remote memories being equally affected by the lesion.

## Methods

### Subjects

Subjects were 17 rhesus macaque monkeys of both sexes (*Macaca mulatta*; mean age 3.9 years, range 2.7-5.1 years, mean weight at time of surgery or equivalent for unoperated controls 4.7 kg, range 3.3-7.4 kg). 4 male monkeys received bilateral neurotoxic hippocampal lesions as described below. The other monkeys (3 female, 10 male) acted as unoperated controls. These experiments were carried out under either the authority of personal and project licences consistent with the United Kingdom Animals (Scientific Procedures) Act 1986, or a protocol approved by the Institutional Animal Care and Use Committee of the Icahn School of Medicine at Mount Sinai. The four monkeys with neurotoxic hippocampal lesions and four of the controls (all male) were tested at Mount Sinai; the remaining monkeys were tested at Oxford University. Data from some of the control monkeys has appeared in previous publications (Mitchell et al., 2008).

### Hippocampal lesions

Monkeys received MRI-guided bilateral neurotoxic hippocampal lesions using methods described by (Hampton et al., 2004). Neurosurgical procedures were performed in a dedicated operating theatre under aseptic conditions. Briefly, monkeys were sedated with a cocktail of dexmedetomidine (0.01 mg/kg), buprenorphine (0.01 mg/kg) and midazolam (0.1 mg/kg) given i.m.. Where necessary, top-ups were given of dexmedetomidine (0.003 mg/kg) and midazolam (0.1 mg/kg) without buprenorphine (to avoid excessive respiratory depression) and any further top-ups of dexmedetomidine (0.003 mg/kg) only as necessary. This protocol was selected to avoid the use of the NMDA antagonist ketamine, which would potentially counteract the effects of the NMDA used as an excitotoxin (Hampton et al., 2004).

Monkeys were intubated, an i.v. catheter placed and anesthesia was maintained with sevoflurane (1.5-4%, to effect, in 100% oxygen). Monkeys were given glycopyrrolate (0.01 mg/kg i.m.), antibiotics (Cefazolin, 25 mg/kg i.m.), steroids (methylprednisolone, 20 mg/kg i.v.), non-steroidal anti-inflammatories (meloxicam, 0.2 mg/kg i.v.), and a H2 receptor antagonist (ranitidine, 1 mg/kg, i.v.) to prevent against gastric ulceration following the administration of both steroids and non-steroidal anti-inflammatories. Atipamezole was used to antagonize the alpha 2-adrenergic agonist if necessary, once anesthesia was stabilized. Monkeys received i.v. fluids throughout the procedure (5 ml/kg/hr i.v.).

The monkey was placed in a stereotaxic frame in exactly the same position as for the preoperative structural MRI scan (employing a tooth marker; (Saunders et al., 1990)). The head was cleaned with antimicrobial cleaner and the skin and underlying galea were opened in layers. Small holes were drilled over the injection entry points: one dorsal and posterior to the long axis of the hippocampus and one dorsal to the uncus in each hemisphere (see Hampton et al., 2004 for details). Two micromanipulators (Kopf Instruments, Tujunga, CA) were fitted with gas-tight syringes (Hamilton, Reno, NV) with a 28 ga needle, point style 4, using measurements obtained from the preoperative T1-weighted scan at the most anterior extent of the hippocampus and injections of N-methyl D-aspartate (NMDA; 0.3 M in sterile saline) were made from anterior to posterior, spaced 1.5 mm apart. Each injection was 1.5-2 µl in volume, made at a rate of 0.5 µl/min, with 1 min between targets. After the final injection the needle was raised 0.5 mm and 10 min elapsed before it was extracted. For the uncus injections 2 injections per hemisphere were made, 3 µl in volume, made at a rate of 0.5 µl/min, with 3 min between targets. Propanolol (0.5 ml of 1 mg/ml per dose) was administered immediately prior to the NMDA injections and re-administered as necessary (up to 4 times) to prevent tachycardia during the injections due to nonspecific effects of NMDA. One monkey received propofol during one surgery (4.0 ml total in boluses of 0.5-1.0 ml of a 10 mg/ml solution) to supplement anesthesia, due to tachypnoea, also likely to be a nonspecific effect of NMDA. Once the lesion was completed the skin and galea were sewn in layers.

When the lesion was complete, monkeys received 0.2 mg/kg metoclopramide (i.m.) to prevent postoperative vomiting. Monkeys also received 0.1 mg/kg midazolam (i.m.) to prevent seizures. They were extubated when a swallowing reflex was evident, returned to the home cage, and monitored continuously until normal posture was regained. Post-operatively monkeys were treated with antibiotics, steroids and analgesia for 3-5 days. Operated monkeys were returned to their social groups within 3 days of the surgery.

Following the first surgery we assessed the lesion extent in each monkey with a T2-weighted scan (Málková et al., 2001) and used the result to plan a second surgery, targeting the injection co-ordinates to regions with low hypersignal. All monkeys received two surgeries.

### Histology

At the end of the study, monkeys were deeply anaesthetized with ketamine (10 mg/kg), intubated and given sodium barbiturate (sodium pentobarbital, 100 mg/kg) intravenously. They were then transcardially perfused with 0.9% saline followed by 4% paraformaldehyde. Brains were post-fixed in paraformaldehyde overnight and then cryoprotected in 30% sucrose solution in 0.9% saline and cut into 50 m sections coronally on a freezing microtome. 1 in 5 sections was stained with cresyl violet for cell bodies. The sections containing the hippocampus were photographed using a Nikon Eclipse 80i light microscope with a 4x objective (Figure 1).

**Figure 1.**
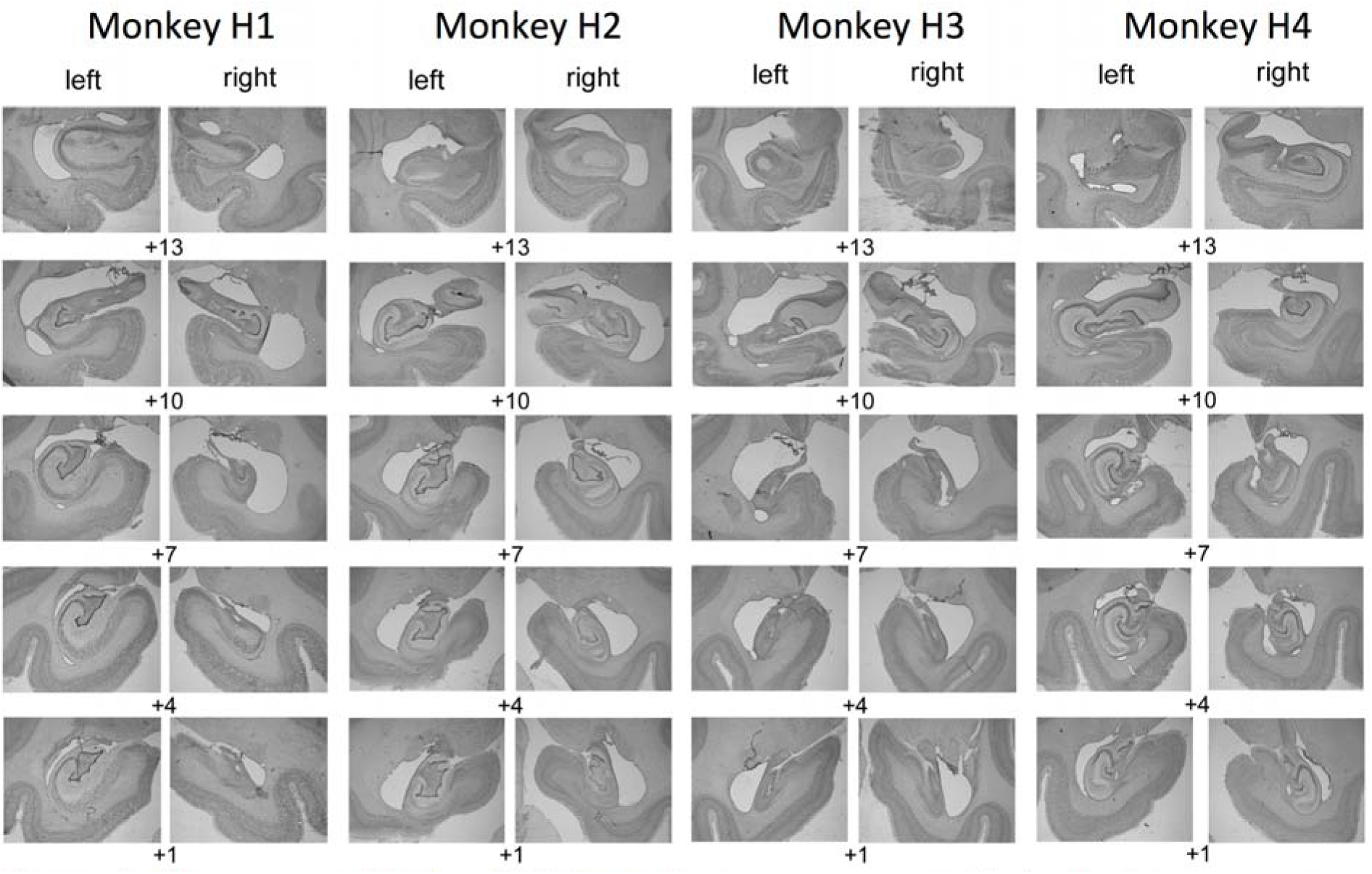
Hippocampal lesion histology. Lesions were specific to the hippocampus in both hemispheres. Cresyl violet stained sections showing the extent of hippocampal lesions in four operated monkeys.

Hippocampal volumetric reduction was carried out in Fiji (https://fiji.sc/), a version of the image analysis program ImageJ. The volume of the hippocampus was manually delineated on sections of the monkey atlas “Red” (using criteria from Málková et al., 2001) and the remaining hippocampal volume of the hippocampus was manually delineated on images of the cresyl violet sections. The sections were then nonlinearly warped to the atlas using the function bUnwarpJ and the volume of each hippocampal section calculated as a percentage of normal hippocampal volume (**Table 1**). The overlap between the remaining hippocampal volume across all 4 monkeys and normal hippocampal volume is shown in **Figure 2**.

**Figure 2.**
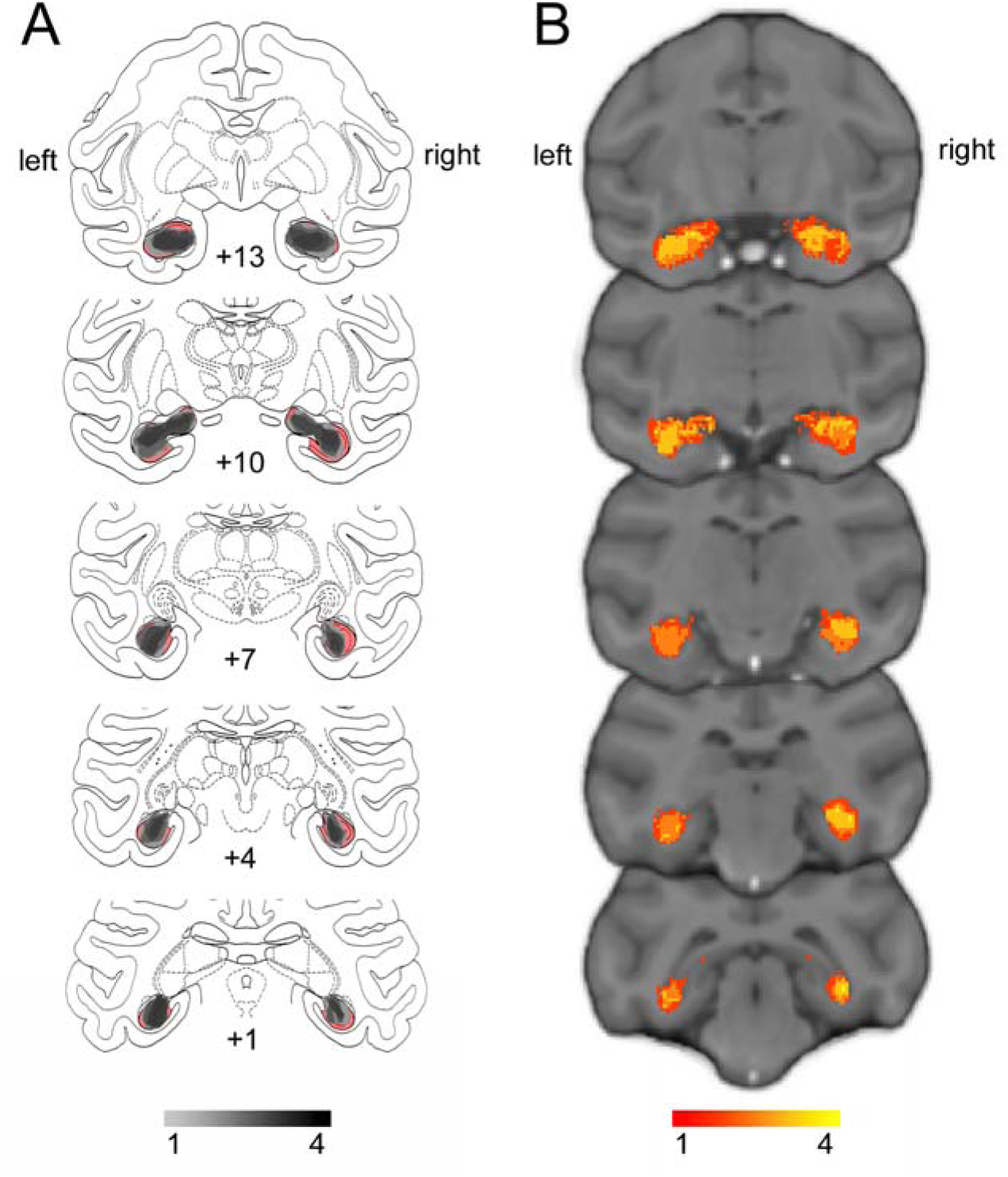
Extent and variability of hippocampal lesions. Sketch of hippocampal size based on histology (Nissl stained sections) overlaid on atlas sections. The unlesioned hippocampal volume is shown in red. Overlap of remaining hippocampal volume is shown for the 4 monkeys indicating shrinkage of the hippocampus bilaterally in all monkeys. **(B)** T2- weighted hypersignal 6 days after surgery indicating local inflammation in the hippocampus; overlap is shown for the 4 monkeys.

**Table 1.** Lesion volumes calculated from Nissl-stained histological sections registered to atlas sections and calculated relative to atlas volumes.

|  | Hippocampus |  |  | Left |  |  | Right |  |  |
| --- | --- | --- | --- | --- | --- | --- | --- | --- | --- |
| Monkey | Normal<br>volume | Lesion | % | Normal<br>volume | Lesion | % | Normal<br>volume | Lesion | % |
| M | 107261.6 | 55354 | **50.93** | 53576.4 | 21646.2 | **37.33** | 53685.2 | 33707.8 | **64.53** |
| N |  | 39578.4 | **37.79** |  | 18372.6 | **33.56** |  | 21205.8 | **42.02** |
| S |  | 66925.6 | **66.02** |  | 32763.8 | **64.01** |  | 34161.8 | **68.03** |
| T |  | 44675.8 | **42.58** |  | 17454.4 | **34.71** |  | 27221.4 | **50.45** |

### Assessment apparatus

Monkeys were assessed using a touchscreen computer for display of stimuli and a linked pellet dispenser for delivery of rewards. At the completion of each correct trial, a 190mg flavored pellet was dispensed into a food cup located below the touchscreen within reach of the monkey. The apparatus also contained a sealed metal lunch box, which automatically opened upon completion of the final trial of the day. For further details see Mitchell et al. (2007).

### Behavioral task

Monkeys were assessed on the object-in-place scenes task. In this task, two alphanumeric characters were shown in the foreground of a scene. In the background of the scene, was a large alphanumeric character, and a number of differently colored and oriented ellipses. The colours of the foreground and background objects were constrained to ensure visibility of the foreground objects via a minimum color difference between each foreground object, and each other object on the screen. One of the two foreground characters was associated with reward (a single pellet). Monkeys had to learn to respond by touching the character that was associated with reward.

### Training

Monkeys were initially trained to touch foreground objects and avoid touching background objects. Following pre-training, monkeys were trained on 25 sequentially presented scenes in a single testing session. This continued for at least 10 sessions. Next the monkeys were given 50 sequentially presented scenes and tested for a minimum of 10 sessions. Following this stage, monkeys were given 100 sequentially presented scenes in a single session. Training continued until monkeys reached a predefined criterion of 90% performance on two consecutive days of testing. Following this stage, all testing sessions involved 100 scene discrimination problems. Monkeys were sequentially tested on three sets of 100 scene discrimination problems, such that a minimum performance of 90% on two consecutive days of testing was required on a set before moving on to the next set of scenes. A touch to the correct object resulted the object flashing for two seconds, and the delivery of a flavored pellet into the cup. A touch to the incorrect object led to the screen immediately turning black, and an increased intertrial interval of 10s. In this case a correction trial was administered, such that the monkey was represented with the scene, but with only the correct option present. In contrast, touches to the scene background resulted in the screen turning black, and the trial being restarted.

Monkeys were tested pre-operatively on the three sets of 100 scene discrimination problems on three consecutive days, on average 82, 46 and 13 days following the last day of training on each set of scenes, respectively.

Following the final hippocampal surgery, monkeys were given 14 days recovery before testing resumed. On the first post-operation testing day monkeys were shown single objects on the screen and rewarded for touching them, in order to become reacquainted with the testing apparatus. On each of the following three days (on average 113, 78 and 45 days following the last day of training on each set of scenes, respectively, and on average 31 days following the pre-operative test), tests on one of the sets of 100 scene discrimination problems occurred, as in the pre-operative testing phase.

### Statistical analysis

The retrograde behavioural data were analysed using a repeated-measures ANOVA, with group, time and set were fixed effects and individual monkeys treated as random effects. For the post-operative new scene learning, a t-test (not assuming equal variances) was used to assess the effect of lesion on the number of errors that occured during learning (until criterion was reached).

## Results

Neurotoxic hippocampal lesions impaired retention of preoperatively acquired memories, irrespective of the time before surgery the memories were learned. Errors made in pre- and postoperative single-trial retention tests, carried out in identical fashion, showed a substantial impact of surgery in monkeys with hippocampal damage, whereas retention was similar in controls tested twice separated by a period of rest. Repeated measures ANOVA on errors in retention test revealed significant main effects of group (F_(1, 15)_ = 9.99, p = .006), pre/postop test (F_(1, 15)_ = 28.14, p < .0005), and, critically, their interaction (F_(1, 15)_ = 8.894, p = .009, **Figure 3A**). There was a main effect of problem set, reflecting better retention of scenes learned closer to the time of surgery (F_(2, 15)_ = 31.17, p < .0005), but this effect did not interact with any others (Fs_(2, 30)_ < 1.67, ps > .21).

**Figure 3.**
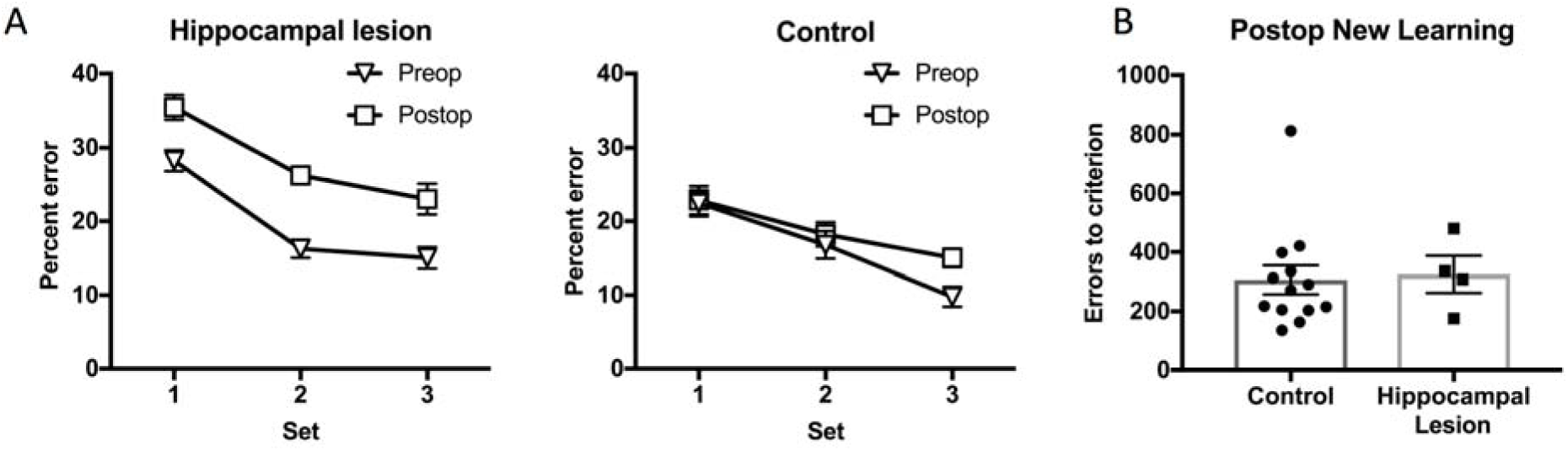
Effects of hippocampal lesions on object-in-place scene memory. A) Retrograde memory impairment in monkeys with hippocampal lesions. Left: Monkeys made a higher proportion of errors on recognition of previously learned object-in-place scenes following hippocampal lesions. Stimulus sets 1-3, learned at different pre-lesion timepoints were equally affected by the lesion. Right: This effect significantly differed from unoperated control monkeys, tested at equivalent timepoints, in whom no effect of (pre- vs post-op equivalent) time was found. B) No anterograde memory impairment in monkeys with hippocampal lesions. Hippocampal lesions caused no discernable impairment in the learning of new scenes, as seen through the number of errors committed en route to reaching the pre-set criterion.

Errors to criterion in new learning of a set of another 100 scene problems were not significantly different between control monkeys and monkeys with neurotoxic hippocampal lesions (t(~7.08) = .235, p = .821, **Figure 3B**).

## Discussion

Rhesus monkeys with bilateral neurotoxic lesions limited to the hippocampus were impaired in retrieval of object-in-place scene problems learned prior to the lesion, but could learn a new set of scene problems at the same rate as controls. Thus, for complex visual scenes, the primate hippocampus is necessary for retrieval, but not new learning. This is congruent with a role for the hippocampus in consolidation of visual memories, to the extent that retrieval of scene problems learned months prior to the hippocampal lesion was impaired. Perhaps surprisingly, there was no indication of any gradient of retrograde amnesia, with equivalent impairment of scenes learned at each time point trained before surgery. The experience of the monkeys with the scenes task and stimuli also did not appear to confer any “schematization” on their memories that rendered them hippocampal-independent.

With regard to the absence of a temporal gradient, it is important to note that we did not explicitly control time before surgery as an experimental variable, with each monkey moving on to the next set of scene problems once the current set was learned to criterion. In practice, acquisition of the most remote set began about four months before surgery and the most recent set a month before surgery. This spans a time interval over which object discrimination problems have been reported to become hippocampal-independent in macaque monkeys (Zola-Morgan and Squire, 1990). Considerable work has been directed at the question of whether the hippocampus plays a time-limited role in memory storage. In general following hippocampal damage or inactivation, studies in rodents have found temporally graded retrograde amnesia of stimulus-stimulus associations in a social transmission of food preference task and temporally extensive retrograde amnesia in spatial learning, both accompanied by anterograde impairments, but mixed results in contextual fear conditioning (Maren et al., 1997; Anagnostaras et al., 1999; Winocur et al., 2013b, reviewed in Winocur et al., 2013a). In this context, a temporally extensive retrograde amnesia for scene problems accompanied by normal anterograde new learning is unusual. An important limitation to any conclusions about a retrograde gradient or lack thereof is that we carried out single-trial retention tests for all scenes prior to surgery. This allowed for a direct comparison of pre- and postoperative retention, but also may have rendered all of the scene problems "recent" and subject to reconsolidation/re-encoding processes that would have re-engaged the hippocampus. This issue bears direct investigation in future studies, perhaps by excluding some scene problems from preoperative retention tests in order to compare effects of lesions or temporary inactivations on “reactivated” scenes versus scenes that had not been tested since learning was complete.

The effect of hippocampal lesions on retention and new learning of object-in-place scene problems in this study was qualitatively and quantitatively almost identical to that of ablations of the anterior entorhinal cortex in an earlier study (Mitchell et al., 2008). This suggests that retrieval of visual scene memories occurs via cortico-cortical interactions between the hippocampus and entorhinal cortex. Supporting this conclusion is the observation that transection of the fornix is without effect on retrieval of preoperatively learned scenes in our paradigm (unpublished data). It also supports the view that cortical and subcortical structures, broadly speaking, have distinct roles in memory acquisition and retrieval (Mitchell et al., 2008; Baxter, 2013).

New learning of scene problems was unimpaired in the monkeys with neurotoxic hippocampal lesions. As with other cortical structures in the primate brain (Thornton et al., 1997; Mitchell et al., 2008), there appears to be a substantial capacity for intact, remaining cortical regions to acquire new memories after focal damage. This may relate to the development of alternative behavioral strategies for new learning after focal brain damage (Manns and Squire, 1999; Squire, 2004), the plastic reorganization of brain networks for memory acquisition (Croxson et al., 2012), or both. We did not ascertain whether newly acquired scene memories after hippocampal damage were as enduring as those in monkeys with an intact hippocampus (Zelikowsky et al., 2012). It seems clear that there are some kinds of representations for which the hippocampus is obligatory, for example those including conjunctions of spatial and temporal information (Bird and Burgess, 2008; Howard and Eichenbaum, 2013), just as other cortical areas are obligatory for other kinds of representations. However, in the intact brain, the hippocampus, like other cortical areas, is engaged in acquisition and storage of visual memories even if its role in representing those memories can be taken over by other cortical areas after damage, supported by the similarity of the present data with neurotoxic hippocampal lesions to effects of entorhinal cortex damage on scene retrieval and new learning (Mitchell et al., 2008), or of rhinal cortex lesions on retrieval and new learning of object discrimination problems (Thornton et al., 1997).

On a broader level, our findings suggest that paradigms for studying memory consolidation and the neural mechanisms of systems consolidation would benefit from extending beyond the investigation of single (or very few) memories that are acquired in a small number of trials. This is obviously beneficial for mechanistic studies in which the time of memory acquisition needs to be precisely known, as for tracking time-dependent biochemical cascades. However, the dynamics of large numbers of unique visual memories, acquired concurrently over a period of time, may engage different mechanisms and may place different demands on different brain structures, or networks of brain structures, as a function of the age of the memory.

## Acknowledgments

Grant support: Wellcome Trust Senior Research Fellowship 071291, Friedman Brain Institute at the Icahn School of Medicine at Mount Sinai.

We thank Peter Sergo for help testing the monkeys, James Young, Charles Adapoe, Frank Macaluso, Ronald Primm, Ignacio Medel, Pedro Hernandez and Lazar Fleysher for help with MRI scans, Bill Janssen for help with perfusions, Richard Saunders for advice on the hippocampal neurosurgical approach, and David Gaffan for advice on experimental design.

## References

Anagnostaras SG, Maren S, Fanselow MS (1999) Temporally graded retrograde amnesia of contextual fear after hippocampal damage in rats: within-subjects examination. J Neurosci 19:1106–1114.

Augustinack JC, van der Kouwe AJ, Salat DH, Benner T, Stevens AA, Annese J, Fischl B, Frosch MP, Corkin S (2014) HM’s contributions to neuroscience: a review and autopsy studies. Hippocampus 24:1267–1286.

Bartsch T, Deuschl G (2010) Transient global amnesia: functional anatomy and clinical implications. Lancet Neurol 9:205–214.

Bartsch T, Döhring J, Rohr A, Jansen O, Deuschl G (2011) CA1 neurons in the human hippocampus are critical for autobiographical memory, mental time travel, and autonoetic consciousness. Proc Natl Acad Sci 108:17562–17567.

Baxter MG (2013) Mediodorsal thalamus and cognition in non-human primates. Front Syst Neurosci 7.

Bayley PJ, Gold JJ, Hopkins RO, Squire LR (2005) The neuroanatomy of remote memory. Neuron 46:799–810.

Bird CM, Burgess N (2008) The hippocampus and memory: insights from spatial processing. Nat Rev Neurosci 9:182.

Cowansage KK, Shuman T, Dillingham BC, Chang A, Golshani P, Mayford M (2014) Direct Reactivation of a Coherent Neocortical Memory of Context. Neuron 84:432–441.

Croxson PL, Browning PGF, Gaffan D, Baxter MG (2012) Acetylcholine Facilitates Recovery of Episodic Memory after Brain Damage. J Neurosci 32:13787–13795.

Ding HK, Teixeira CM, Frankland PW (2008) Inactivation of the anterior cingulate cortex blocks expression of remote, but not recent, conditioned taste aversion memory. Learn Mem 15:290–293.

Frankland PW, Bontempi B, Talton LE, Kaczmarek L, Silva AJ (2004) The involvement of the anterior cingulate cortex in remote contextual fear memory. Science 304:881–883.

Frankland PW, Ding H-K, Takahashi E, Suzuki A, Kida S, Silva AJ (2006) Stability of recent and remote contextual fear memory. Learn Mem 13:451–457.

Gaffan D (1994) Scene-specific memory for objects: a model of episodic memory impairment in monkeys with fornix transection. J Cogn Neurosci 6:305–320.

Goshen I, Brodsky M, Prakash R, Wallace J, Gradinaru V, Ramakrishnan C, Deisseroth K (2011) Dynamics of retrieval strategies for remote memories. Cell 147:678–689.

Hampton RR, Hampstead BM, Murray EA (2004) Selective hippocampal damage in rhesus monkeys impairs spatial memory in an open-field test. Hippocampus 14:808–818.

Howard MW, Eichenbaum H (2013) The hippocampus, time, and memory across scales. J Exp Psychol Gen 142:1211.

Kapur N, Brooks DJ (1999) Temporally-specific retrograde amnesia in two cases of discrete bilateral hippocampal pathology. Hippocampus 9:247–254.

Málková L, Lex CK, Mishkin M, Saunders RC (2001) MRI-based evaluation of locus and extent of neurotoxic lesions in monkeys. Hippocampus 11:361–370.

Manns JR, Squire LR (1999) Impaired recognition memory on the Doors and People Test after damage limited to the hippocampal region. Hippocampus 9:495–499.

Maren S, Aharonov G, Fanselow MS (1997) Neurotoxic lesions of the dorsal hippocampus and Pavlovian fear conditioning in rats. Behav Brain Res 88:261–274.

Mitchell AS, Browning PG, Wilson CR, Baxter MG, Gaffan D (2008) Dissociable roles for cortical and subcortical structures in memory retrieval and acquisition. J Neurosci 28:8387–8396.

Moscovitch M, Rosenbaum RS, Gilboa A, Addis DR, Westmacott R, Grady C, McAndrews MP, Levine B, Black S, Winocur G (2005) Functional neuroanatomy of remote episodic, semantic and spatial memory: a unified account based on multiple trace theory. J Anat 207:35–66.

Reed JM, Squire LR (1998) Retrograde amnesia for facts and events: findings from four new cases. J Neurosci 18:3943–3954.

Saunders RC, Aigner TG, Frank JA (1990) Magnetic resonance imaging of the rhesus monkey brain: use for stereotactic neurosurgery. Exp Brain Res 81:443–446.

Scoville WB, Milner B (1957) Loss of recent memory after bilateral hippocampal lesions. J Neurol Neurosurg Psychiatry 20:11.

Squire LR (2004) Memory systems of the brain: a brief history and current perspective. Neurobiol Learn Mem 82:171–177.

Tanaka KZ, Pevzner A, Hamidi AB, Nakazawa Y, Graham J, Wiltgen BJ (2014) Cortical representations are reinstated by the hippocampus during memory retrieval. Neuron 84:347–354.

Tayler KK, Tanaka KZ, Reijmers LG, Wiltgen BJ (2013) Reactivation of Neural Ensembles during the Retrieval of Recent and Remote Memory. Curr Biol 23:99–106.

Teixeira CM, Pomedli SR, Maei HR, Kee N, Frankland PW (2006) Involvement of the Anterior Cingulate Cortex in the Expression of Remote Spatial Memory. J Neurosci 26:7555–7564.

Thornton JA, Rothblat LA, Murray EA (1997) Rhinal cortex removal produces amnesia for preoperatively learned discrimination problems but fails to disrupt postoperative acquisition and retention in rhesus monkeys. J Neurosci 17:8536–8549.

Tse D, Takeuchi T, Kakeyama M, Kajii Y, Okuno H, Tohyama C, Bito H, Morris RGM (2011) Schema-Dependent Gene Activation and Memory Encoding in Neocortex. Science 333:891–895.

Winocur G, Moscovitch M, Sekeres MJ (2013a) Factors affecting graded and ungraded memory loss following hippocampal lesions. Neurobiol Learn Mem 106:351–364.

Winocur G, Sekeres MJ, Binns MA, Moscovitch M (2013b) Hippocampal lesions produce both nongraded and temporally graded retrograde amnesia in the same rat. Hippocampus 23:330–341.

Zelikowsky M, Bissiere S, Fanselow MS (2012) Contextual fear memories formed in the absence of the dorsal hippocampus decay across time. J Neurosci 32:3393–3397.

Zola-Morgan SM, Squire LR (1990) The primate hippocampal formation: evidence for a time-limited role in memory storage. Science 250:288–290.

